# Analysis of Hippocampal Synaptic Function in a Rodent Model of Early Life Stress

**DOI:** 10.1101/2024.04.29.591711

**Authors:** Matthew P Wilkinson, Emma S J Robinson, Jack R Mellor

## Abstract

**Background:** Early life stress (ELS) is an important risk factor in the aetiology of depression. Developmental glucocorticoid exposure impacts multiple brain regions with the hippocampus being particularly vulnerable. Hippocampal mediated behaviours are dependent upon the ability of neurones to undergo long-term potentiation (LTP), an N-methyl-D-aspartate receptor (NMDAR) mediated process. In this study we investigated the effect of ELS upon hippocampal NMDAR function.

**Methods:** Hooded Long-Evans rat pups (n=82) were either undisturbed or maternally separated for 180 minutes per day (MS180) between post-natal day (PND) 1 and PND14. Model validation consisted of sucrose preference (n=18) and novelty supressed feeding (NSFT, n=34) tests alongside assessment of corticosterone (CORT) and paraventricular nucleus (PVN) cFos reactivity to stress and hippocampal neurogenesis (all n=18). AMPA/NMDA ratios (n=19), miniEPSC currents (n=19) and LTP (n=15) were assessed in whole-cell patch clamp experiments in CA1 pyramidal neurones.

**Results:** MS180 animals showed increased feeding latency in the NSFT alongside increased overall CORT in the restraint stress experiment and increased PVN cFos expression in males but no changes in neurogenesis or sucrose preference. MS180 was associated with a lower AMPA/NMDA ratio with no change in miniEPSC amplitude or area. There was no difference in short- or long-term potentiation between MS180 and control animals nor were there any changes during the induction protocol.

**Conclusions:** The MS180 model showed a behavioural phenotype consistent with previous work. MS180 animals showed increased NMDAR function with preliminary evidence suggesting that this was not concurrent with an increase in LTP.

## 1. Introduction

Early life stress (ELS) predisposes to multiple neuropsychiatric disorders including depression, anxiety and schizophrenia (Agid et al., 1999; Green et al., 2010; Lemoult et al., 2019; McCauley et al., 1997). Events during childhood that exceed a child’s ability to cope generate elevated levels of glucocorticoids. This leads to an altered neurodevelopmental trajectory in areas of the brain that have high glucocorticoid receptor (GR) expression and a slow developmental trajectory such as the prefrontal cortex (PFC), amygdala and hippocampus (Cohodes et al., 2020; Tottenham and Sheridan, 2009). However, how ELS leads to an increased risk of developing psychiatric disorders is still poorly understood (Targum and Nemeroff, 2019).

Reward processing deficits have been suggested to be a key feature of both depression (Admon and Pizzagalli, 2015; Halahakoon et al., 2020; Pizzagalli et al., 2005) and ELS (Novick et al., 2018; Pechtel and Pizzagalli, 2011, 2013; Wilkinson et al., 2021) with the potential to form an intermediate phenotype. The hippocampus is both susceptible to glucocorticoids and a key region involved in the processing of reward (Dupret et al., 2013; Hok et al., 2007; Rolls and Xiang, 2005; Shohamy et al., 2009) therefore may mediate some of the effects of ELS. One of the ways in that hippocampal information processing may be altered by ELS is through changes in N-methyl-d-aspartate receptor (NMDAR) function with NMDARs being crucial for the long-term potentiation (LTP) that is believed to underly learning and memory (Collingridge and Bliss, 1987; Volianskis et al., 2013). It is well understood how acute glucocorticoid exposure can alter NMDAR function (Mikasova et al., 2017; Tse et al., 2012; Xiao et al., 2010), however the impact of long term exposure during development is poorly understood. A previous meta-analysis investigated the effects of ELS upon LTP in hippocampal CA1 using rat models of ELS (Derks et al., 2017). LTP was found to be increased in maternally separated rats, the most common animal model of ELS, while lower in a different model using natural variations in maternal care. Other studies, not included in this meta-analysis, have also reported lower LTP in maternally separated animals compared to controls (Cao et al., 2014; Heydari et al., 2019; Sousa et al., 2014). Previous reports have suggested a decreased NMDAR relative to AMPAR function following ELS (Pillai et al., 2018) in addition to decreased GluN2B subunit expression (Lesuis et al., 2019; Pickering et al., 2006; Roceri et al., 2002). Another model of psychiatric risk, DLG2 heterozygous deletion, was recently associated with impaired associative LTP but increased NMDAR function due to increased potassium leak channel function (Griesius et al., 2022) alongside a mild reward processing impairment (Griesius et al., 2023).

However, a major drawback of previous work is that it has not considered the key factors of animal sex, hippocampal region, and input pathway. There is evidence that females are more susceptible to the effects of ELS in humans while the opposite is true in rodent models (Bonapersona et al., 2019; Herbison et al., 2017). Additionally there are changes in LTP induction rules between sexes (Qi et al., 2016; Tozzi et al., 2019; Warren et al., 1995; Wei et al., 2018). The dorsal (DH) and ventral (VH) hippocampus should also be treated separately with the ventral hippocampus being more associated with reward learning and the dorsal hippocampus being responsible more for spatial learning (Fanselow and Dong, 2010). There are differences between LTP induction rules between the DH and VH with changes in SK channel regulation of synaptic excitability and dendritic plateau potentials between the regions (Babiec et al., 2017; Kouvaros and Papatheodoropoulos, 2016; Malik and Johnston, 2017). Finally, the role of Temporoammonic (TA) and Schafer collateral (SC) pathways in the hippocampus have never been appreciated with respect to ELS.

To consider hippocampal LTP, NMDAR function and circuit dynamics following ELS in a rigorous manner it is critical to consider all these factors which likely interact in complex relationships to lead to behavioural changes and neuropsychiatric risk in both rodents and humans. In this study we bred a cohort of rats using the maternal separation model of ELS (Mirescu et al., 2004) then validated successful application of the model using a battery of behavioural and biochemical assays matching previous work (Stuart et al., 2019). We then conducted *ex-vivo* patch clamp electrophysiology in hippocampal CA1 to measure the ratio of AMPAR and NMDAR mediated excitatory post synaptic currents (EPSCs) before then measuring miniature EPSCS (miniEPSCs) to localise any changes in AMPAR/NMDAR ratio to either changes in AMPAR or NMDAR function. We also assessed changes in LTP in MS180 animals, however these experiments were hampered by sample size limitations due to the Covid-19 pandemic.

By understanding how early life stress influences hippocampal CA1 circuit dynamics this may allow a greater understanding into the links between ELS, reward learning deficits and the development of psychiatric disease. Further understanding into these effects may also allow the identification of novel targets that may either be beneficial in treating depression directly or reducing the risk to those who have suffered elevated levels of stress in childhood from developing mental health conditions.

## 2 Methods

### 2.1 Animals

A total of 19 hooded long-Evans rats and their 82 offspring were used with rats being derived from an in-house breeding colony. While breeding, animals were housed in standard lighting conditions (12:12h cycle, lights off at 19:00) with offspring used for electrophysiology remaining in these conditions. Animals used for validation were transferred to reverse lighting conditions (12:12h cycle, lights on at 20:30) at least 2 weeks before the start of experiments. Animals had free access to food and water and were provided with wooden chew blocks and red houses as enrichment in temperature and humidity controlled conditions. Sample size was estimated from previous studies (Stuart et al., 2019). All experiments were undertaken in accordance with local institutional guidelines, the UK Animals (Scientific procedures) Act of 1986 and the European Community Council Directive of 24 November 1986 (86/609/EEC).

### 2.2 Study Design

MS180 animals were bred in two separate cohorts (see figure 1 for an overview of the entire study). Cohort 1 animals were subdivided into two groups following weaning: an electrophysiology and a validation group. The validation group completed the novelty supressed feeding test (NSFT), sucrose preference and restraint stress corticosterone experiments before being terminally used in restraint stress cFos and BrdU neurogenesis experiments. The electrophysiology group from cohort 1 were used for AMPA/NMDA ratio and miniEPSC experiments. Animals from cohort 2 were used to provide additional power in the NSFT experiment before some were used for AMPA/NMDA and miniEPSC experiments (5 rats from 2 litters) with the remainder being used in LTP experiments. During all experiments following animal weaning the experimenter was blind to animal treatment and all efforts were made to counterbalance for sex and animal condition in experiments. In electrophysiology studies further distinction was made between studying both hippocampal region and CA1 input pathway with this described in detail later. Four other breeding cycles were also initiated with 2 incidences of dams failing to get pregnant, one incidence of litters being rejected by the dam and one incident of animals having to be culled due to Covid-19.

**Figure 1.**
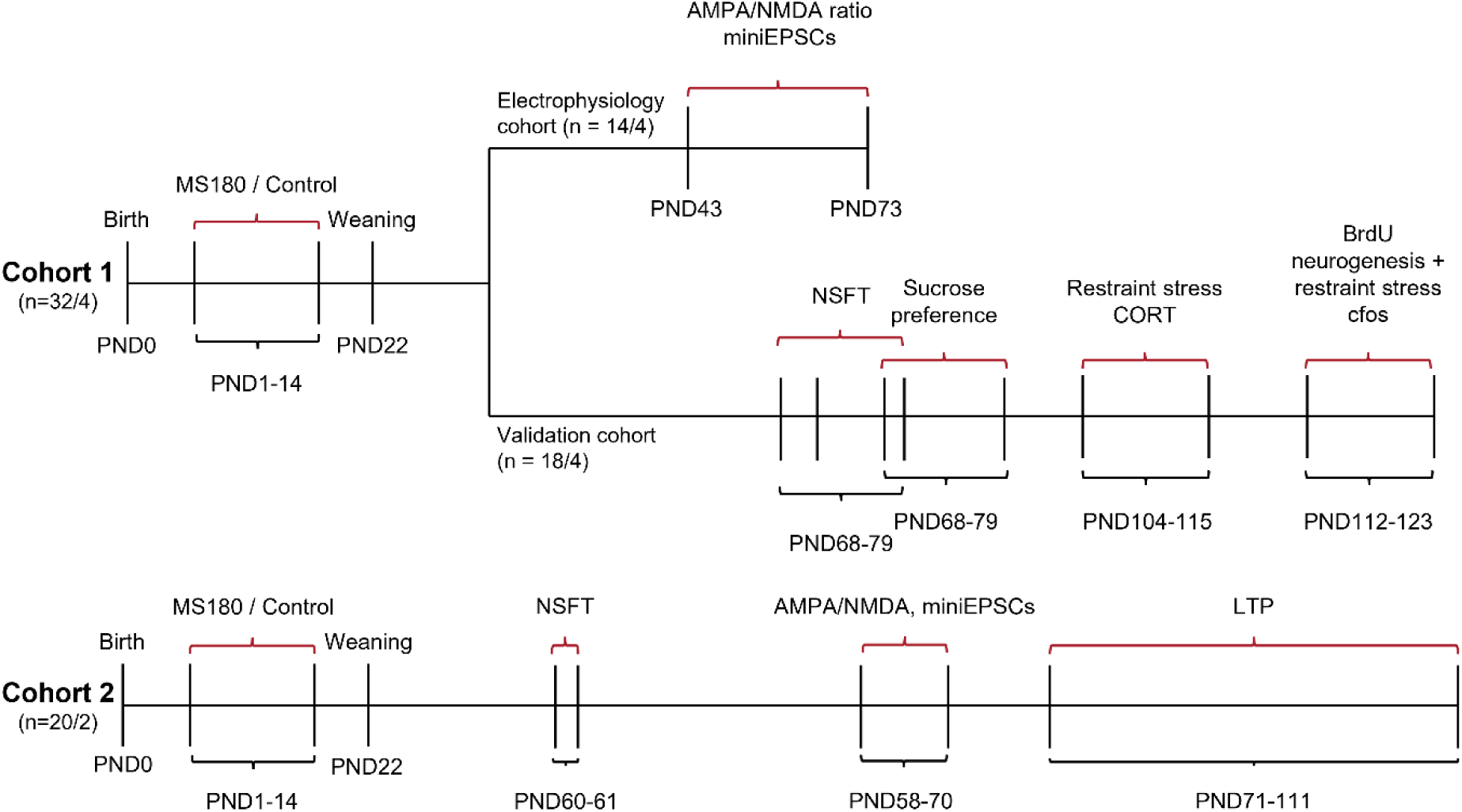
Study overview. Animals were generated in breeding cycles with animals being allocated from these into different experiments. n = 32/4 refers to 32 animals from 4 litters.

### 2.3 Maternal separation procedure

Maternal separation procedures were completed as described by Mirescu et al., 2004. Male and female rats were either pair housed or housed as trios (1 male and 2 females) for 2 weeks or until females were visibly pregnant. Pregnant animals were then singly housed and monitored daily for signs of birth. Animals were born on PND0 before having litter sizes adjusted to 8 pups (Cohort 1) or 10 pups (Cohort 2) on PND1 with equal numbers of male and female animals preferred. Litters were either randomised to control or MS180 conditions on PND1. MS180 pups were subject to daily separations of 180 minutes from PND1 to PND14 away from the dam and were placed into an incubator held at 32°C during this period. Separation always occurred between 13:00 and 16:00 with pups remaining as a litter in a container containing sawdust and bedding from the home cage. Control litters were left completely alone with no interventions. All cage cleaning halted during the 14-day experimental period. Following PND14 all animals were returned to standard husbandry with cage cleaning resuming and animals were weaned at PND22 into groups of 2-3 by sex and by litter. At this point animals from the Cohort 1 validation group were transferred to reverse lighting while animals destined for electrophysiology were kept under standard lighting.

### 2.4 Model validation experiments

#### 2.4.1 Novelty supressed feeding test

Experiments were performed as described by Stuart et al., 2019 during animals’ active phase. Animals were food deprived for 24 hours before being placed into a 70cm diameter arena containing a 10cm food bowl filled with standard chow placed into the centre of the arena. Latency to both approach the bowl and eat from it were recorded and animals were recorded with a Logitech C920 webcam. Up to 15 minutes was allowed for animals to eat from the bowl before they were removed from the arena and classed as failing to eat. Tracking was performed utilising Noldus Ethovision XT software to generate the additional parameters of percentage time moving and average velocity. The area was additionally subdivided into an inner and outer zone with this being used to calculate percentage time in the inner zone.

#### 2.4.2 Sucrose preference test

Animals were first habituated to 1% sucrose in tap water with this being provided via two sipper sacks (Edstrom-Avidity Science, USA) for two days before animals were provided with standard drinking water from sipper sacks for a day. Animals were then water restricted for 4 hours and individually housed in test cages 90 minutes before testing. Animals were then provided with one sipper sack containing 1% sucrose and another containing standard drinking water. Consumption was measured at 30 min, 60 min and 120 min after the start of testing with bottle positions swapped at these timepoints. Following the final timepoint animals returned to their home cages and sucrose preference was calculated.

#### 2.4.3 Restraint stress corticosterone

Animals’ corticosterone response to restraint stress was assessed within-subject through tail vein blood sampling. Animals first had their cage heated on a warming mat in their holding room for 10 minutes before being moved to a procedure room and their tail being directly heated with a warming mat 1 minute before baseline sample acquisition. A baseline sample of approximately 250µl was collected into tube containing 50µl of ice-cold EDTA before animals were placed into a restraint stress tube (Harvard Apparatus, USA) for 20 minutes. Just prior to the end of the 20-minute period animals had their tail heated again before another tail vein blood sample was collected. The experiment took place over two days with animals being allocated to each day such that the study was counterbalanced for condition and sex but the experimenters remained blind to condition. All blood samples were stored on ice before being centrifuged at 8000g for 10 minutes and the plasma being aspirated off before being frozen at −20°C prior to further analysis. Plasma CORT concentrations were assessed using previously described methods (George et al., 2017) with final concentrations being adjusted for the blood volume collected.

#### 2.4.4 cFos and BrdU Immunohistochemistry

Animals were dosed with 50mg/kg BrdU (Sigma, USA) in 0.7% saline i.p every 2 hours four times and 22.5 hours following the final injection were subject to 20 minutes of restraint stress. This happened when animals were between PND 112 and PND 123, a point where neurogenesis is decreasing steeply with age (Merkley et al., 2014) and older than the age where neurogenesis differences between MS180 and control groups were previously observed (Stuart et al., 2019). 24 hours following the final BrdU injection animals were deeply anaesthetised with pentobarbital (Merial, UK) before being trans cardiac perfused with first ice cold phosphate buffer (PB, 28mM NaH_2_PO_4_, 72mM Na_2_HPO_4_) and then ice cold 4% paraformaldehyde (PFA) in PB. Following adequate fixation brains were removed and then placed overnight into 4% PFA in PB. Brains were then transferred into a 25% sucrose solution until they sunk and were then snap frozen in optimal cutting temperature compound (Scigen, USA) using dry ice. Brains were cut into 40µm thick sections using a freezing microtome (Reichert, Austria) before being frozen in cryoprotectant (30% sucrose, 30% ethylene glycol, 50% PB in H_2_O) prior to use in immunohistochemistry (IHC) experiments.

Hippocampal sections (−1.92mm ≤ bregma ≤ −6.48mm) were used for neurogenesis experiments while sections containing PVN (−1.56mm ≤ bregma ≤ 1.92mm) were used for restraint stress cFos experiments. cFos IHC was conducted using Tris-buffered saline (TBS: 100mM Tris, 155mM NaCl, pH 7.4) while BrdU IHC utilised phosphate buffered saline (PBS: 1.45mM KH_2_PO_4_, 8.1mM Na_2_HPO_4_, 136.5mM NaCl, 2.68mM KCl, pH 7.2). All sections were first washed with buffer 4 times for 10 minutes while being subject to gentle agitation. At this point sections in the BrdU experiment completed an additional step consisting of a 30m incubation in 2M HCl at 37°C followed by neutralisation with two 5-minute washes with 0.1M NaB_4_O_7_ followed by another wash phase with buffer (3×5 min). Sections from both experiments were then blocked for non-specific binding for 90 minutes in 2% bovine serum albumin and 3% serum (goat for cFos and donkey for BrdU experiments) in buffer. Primary antibody (cFos: 1:4000 polyclonal rabbit anti-cFos, Merck ABE457, lot numbers 2935662/2967893, BrdU: 1:100 monoclonal mouse anti-BrdU, BD Biosciences B44, lot number 9172603) was then applied overnight at room temperature within 3% serum in buffer. Following another wash phase (3 x 5 min in buffer) sections were then incubated in secondary antibody (cFos: 1:200 polyclonal goat anti-rabbit IgG(H+L) Alexa fluor 594, Thermofisher A11037, lot number 1851471, BrdU: 1:500 Polyclonal donkey anti-mouse IgG(H+L) Alexa fluor 488, Thermofisher A21202, lot number 1696430) diluted in 3% serum in buffer for 2 hours before another wash (3 × 5 min in buffer). Sections were then incubated in a 1:1000 dilution of DAPI for 2 minutes before again being washed (3 × 5m) and mounted on distilled water using vectashield (Vectorlabs, US) fluorescent mounting medium. Images were captured at 10x magnification using a Leica DMI6000 widefield microscope with DFC365FX camera using LASX acquisition software (cFos: 5s @ 6x gain exposure, BrdU: 150ms @ 5x gain). Cell counting was conducted manually using ImageJ. Due to observed ice damage in PVN sections this was quantified on a 10-point scale and used as a random factor in analysis; there was no correlation between ice damage and cFos count (data not shown). For the BrdU experiment sections were counted as being dorsal (bregma ≥ −3.44mm), intermediate (−3.44mm ≥ bregma ≥ −5.08mm) or ventral (bregma ≤ −5.08mm).

### 2.5 Electrophysiology experiments

#### 2.5.1 Slice preparation

Transverse hippocampal slices were prepared from rats following terminal anaesthesia under isoflurane and decapitation. Following removal of the brain hippocampi were dissected in ice cold cutting solution (in mM: Sucrose 205, Glucose 10, NaHCO_3_ 26, KCl 2.5, NaH_2_PO_4_ 1.25, CaCl_2_ 0.5, MgCl_2_ 5) bubbled with 95% O_2_, 5%CO_2_ before being sliced into 400µm transverse slices using a Leica LS1200 vibratome. Brain slices were immediately transferred into aCSF (in mM: NaCl 124, Glucose 10, NaHCO_3_ 24, KCl 3, NaH_2_PO_4_ 1.25, CaCl_2_ 2.5, MgCl_2_ 1.3), again bubbled with 95% O_2_ and 5%CO_2_, and incubated at 35°C for 30 minutes then a further 30 minutes at room temperature prior to the start of patch clamp experiments. Dorsal and ventral slices were classified as those coming from the extreme 1/3 of the hippocampal axis.

#### 2.5.2 Whole cell patch clamp recordings

Prior to transfer into a submerged slice chamber for whole-cell patch clamp experiments all slices had CA3 manually removed. Slices were submerged in a constant flow (≈2.5ml/min) of aCSF and held at 32°C. Slices were stimulated (protocols described later) by tungsten bipolar electrodes (Microprobes for Life Science, USA) and visualised utilising differential interference contrast microscopy using an Olympus BX51WI upright microscope. Patch pipettes with resistance 2-9MΩ were pulled using a Sutter P-97 from borosilicate glass (Harvard apparatus, USA) before being filled with either caesium based (in mM: CsMeSO_3_ 117, HEPES 10, EGTA 0.3, Mg-ATP 2, Na-GTP 0.3, NaCl 9, TEA 10, QX314 1) or potassium based internal solution (in mM: KMeSO_3_ 120, HEPES 10, EGTA 0.2, Mg-ATP 4, Na-GTP 0.3, NaCl 8, KCl 1). Recordings were made using a MultiClamp 700A amplifier (Axon instruments, US) coupled to a CED Micro 1401 digitiser. Data was captured using Signal version 5 (CED, UK. Open access alternative: <u>WinWCP</u>), filtered at 2.4kHz and digitised at 10kHz with either 2x gain (AMPA/NMDA ratio experiments) or 20x gain (miniEPSC experiments). For all experiments series resistance (R_ser_) and input resistance (R_in_) was measured through injection of a 20pA square pulse lasting 500ms with cells being excluded if their R_ser_ increased above 35MΩ. In all experiments junction potential was not corrected for and cells were perfused with 50µM picrotoxin (PTX, Sigma Aldrich, USA).

#### 2.5.4 AMPAR/NMDAR ratio measurement

Cells were held at −70mV in voltage clamp (V_clamp_) for 10 minutes before the start of recording to ensure that LTP would not contaminate recordings. Each pathway was then stimulated in a paired pulse protocol whereby two stimulations separated by 100ms were delivered to generate excitatory post synaptic currents (EPSCs) of approximately 100pA amplitude. Each pathway was stimulated every 10s sequentially such that the SC pathway was stimulated followed 5s later by the TA pathway and then after another 5s the SC pathway was stimulated again. Stimulation was delivered at −70mV for 5 minutes to isolate the AMPA mediated component of the EPSC before cells were held at +40mV to allow NMDAR activity. Following being held at +40mV cells were then returned to −70mV and if greater than a 50% change in EPSC amplitude or R_ser_ was observed then the cell was excluded from analysis. From these recordings an AMPA / NMDA ratio was calculated as:

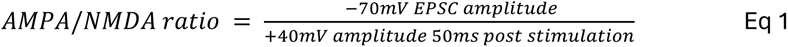

A paired pulse ratio was also calculated from traces as the ratio between the amplitudes of the first and second EPSC in the paired pulse stimulation protocol.

#### 2.5.5 miniEPSC recordings

Neurones were held at −70mV in V_clamp_ while exposed to 500nM tetrodotoxin (TTX, Hello Bio, UK) and were recorded for 5 minutes to record miniEPSC events. Recorded traces were analysed for events using WinEDR v3.9 (Strathclyde university, UK) using a detection template (amplitude: −3pA, tau rise: 0.1ms, tau decay 3ms, dead time: 15ms, rising edge window: 2ms). Detected events were then filtered (10ms ≤ duration ≤ 1000ms, 4ms ≤ tau rise ≤ 1000ms, 0 pAms^-1^ ≤ area ≤ 1000 pAms^-1^) before remaining events were manually screened for inclusion. Inter-event interval was calculated from the per-event averages whereas amplitude and area were calculated from the average miniEPSC trace of each cell.

#### 2.5.6 LTP Experiments

CA1 pyramidal neurones were held at −70mV in V_clamp_ and stimulated through bipolar electrodes placed in the slm (TA pathway), orthodromic sr and antidromic sr to stimulate the SC pathway therefore forming a control and test SC pathway. Cells were stimulated in a paired pulse stimulation protocol to produce a ≈100pA EPSC followed by another stimulation 50ms later in each pathway. Pathways were stimulated sequentially every 5 seconds. For the length of the experiment cells were recorded using the KMeSO_4_ internal. A 5-minute baseline period was recorded before LTP was induced using a theta burst stimulation (TBS) protocol (Buchanan and Mellor, 2007) no more than 10 minutes after a whole cell configuration was achieved to avoid the washout of LTP. To induce LTP, cells were switched to current clamp (I_clamp_) with current injected to maintain the cell at −55mV before three trains of TBS were applied to the test pathways with a 10s interval. The TBS trains consisted of 10 bursts with an inter-burst frequency of 5Hz and each burst containing 5 stimulations at a frequency of 100Hz. Following LTP induction cells were returned to −70mV in V_clamp_ and again stimulated sequentially for another 30 minutes. Responses were normalised in each pathway to the EPSC amplitude during the baseline period and STP was taken to be the response 0-5 min post TBS while LTP was taken to be the period 25-30m post TBS. Cells were excluded from analysis if the series resistance or control pathway amplitude increased by over 50% during a recording. Spikes during the TBS train were counted before being removed to analyse the hyperpolarisation induced by the TBS. Due to the hyperpolarisation following the first theta burst, a baseline was interpolated for each 2s theta burst train consisting of three segments. The area of each theta burst was calculated as the integral of V_m_ with respect to the fitted baseline while the area of the decay phase of each burst was calculated to be the integral of each burst with respect to the fitted baseline following the maximal V_m_ deflection. Paired pulse ratio was calculated from traces as the ratio between the amplitudes of the first and second EPSC in the paired pulse stimulation protocol while input resistance (Rin) was calculated from the end of every trace by the injection of a 20pA square pulse lasting 500ms.

### 2.6 Statistical Analysis

Due to a lack of power due to low n-number in the LTP experiments due to the Covid-19 pandemic two statistical approaches were taken. Model validation experiments, AMPAR/NMDAR ratio and miniEPSC experiments used a multi-level model approach accounting for sex, region, and pathway while LTP experiments were analysed using more traditional ANOVAs and T-tests and did not account for other factors.

#### 2.6.1 Multilevel Analysis

In conducting a complex study consisting of multiple nested factors it is necessary to account for these in the statistical analysis approach in order to avoid pseudoreplication (Lazic et al., 2020). Due to this a multi-level approach was implemented considering the multilevel structure: *Cell* ∈ *Animal* ∈ *Litter*. Due to the extended time-period over which electrophysiology experiments occurred a random factor of age was also included. A generalised linear mixed model (GLMM) was fitted using the glmmTMB package in R version 4.0 (Brooks et al., 2017; R Core Team, 2020). Models were first fit with all possible main factor combinations before being subject to stepwise removal of terms using Akaike information criterion (AIC) for model term deletion. Multilevel and other random effects were always included as a random intercept. Once the simplest model was achieved, the point at which removing terms decreased the model fit, the final model was compared with a null model containing no fixed predictors using AIC calculated from the bbmle package (Bolker and R Development Core Team, 2020). Assuming that the model containing predictors was a better fit to data then the normality and homoscedasticity of residual assumptions were checked using visual methods in addition to Shapiro-Wilk and Breusch-Pagan tests (Zeileis and Hothorn, 2002). Where a satisfactory model fit could not be made using a gaussian error family, efforts were made to both use altered link functions and altered error families. The R^2^ value for each model was also always calculated to assess ultimate model fit (Barton, 2020). Estimated marginal means, the change in outcome where one factor is changed and all other remain constant, for the top level factors of condition, sex, region and pathway were calculated where appropriate using the ggeffects package (Ludecke, 2018). Where at least a trend towards an interaction was observed the this was investigated through fitting of the relevant simplified model where each factor was assessed in turn. For data from PVN cFos and BrdU neurogenesis experiments each section was included in the analysis with an additional random factor of ice damage for the cFos experiment.

#### 2.6.2 Other statistical analysis

For LTP experiments, due to low power resulting from Covid-19 disruption either 2-way ANOVAs or t-tests were used. For LTP analysis 2-way ANOVAS (factors: condition, pathway) were used with post-hoc comparisons being conducted with Dunnet’s multiple comparison test. All data were assessed for normality using Shapiro Wilk and Kolmogorov-Smirnov tests. Data that passed the assumptions of normality was also assessed for violations of Sphericity using Mauchly’s test and where this assumption was violated then the Huynh-Feldt correction was used to adjust the degrees of freedom. Unpaired T-tests were conducted for TBS output measures and input resistance. 2-way ANOVAs and t-test were conducted in GraphPad Prism 10. All graphs were constructed using GraphPad Prism 8 with main effects indicated over the relevant data with a bar and stars. Main effects from the overall mixed model have been indicated on marginal means with the # symbol to emphasise that these are predicted marginal means. All data is shown as mean ± SEM except for marginal means which are mean ± 95% confidence intervals (CI). #/* ≤ 0.05, ##/** < 0.01, ###/*** < 0.001, ####/**** < 0.0001.

## 3 Results

### 3.1 Model Validation

In order to validate that a phenotype mirroring that previously seen in the MS180 model (Stuart et al., 2019) had been generated, animals first completed the novelty supressed feeding test (NSFT). MS180 animals took longer to feed from the bowl than controls (Figure 2A, GLMM, Z = −2.71, p = 0.007) while showing no difference in time to approach the bowl (Figure 2B). When the proportion of time animals spent in the inner zone of the arena compared to the outer zone was analysed a difference between groups emerged (Figure 2C, GLMM, Z = −2.62, p = 0.009) with MS180 animals spending less time in the inner zone compared to controls. Animals additionally completed the sucrose preference test, a measure of reward sensitivity. There was no difference between groups in sucrose preference (Figure 2D) however animals did show an overall sucrose preference (Wilcoxon signed ranks test against hypothetical mean of 50%, Z = 2.98, p = 0.003).

**Figure 2.**
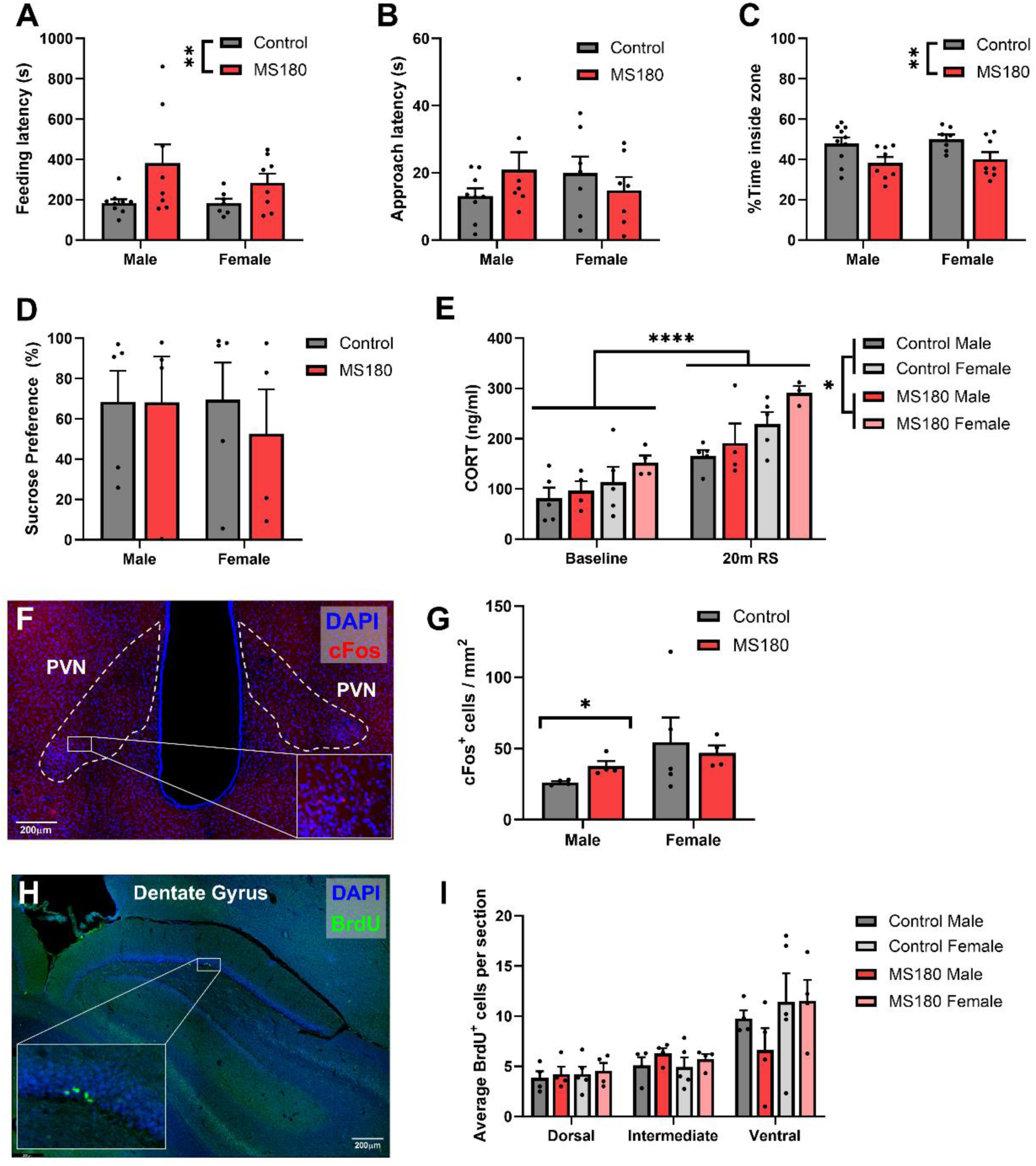
Validation of the MS180 model. **(A-C)** Feeding latency, approach latency and time spent in the centre of the arena from the novelty supressed feeding test. (**D)** Sucrose preference after 120 minutes **(E)** Plasma corticosterone concentration before and after 20 minutes of restraint stress **(F,G)** Example photomicrograph showing cFos expression in the PVN and cFos^+^ cell count after 20 minutes of restraint stress **(H,I)** Example photomicrograph of the dentate gyrus and BrdU^+^ cell count in the subgranular zone indicating levels of neurogenesis. NSFT: n = 33 animals (17 control, 16 MS180); restraint stress CORT and sucrose preference: n = 18 (10 control, 8 MS180); PVN c-Fos and BrdU neurogenesis: n=17 (9 control, 8 MS180). Data shown as mean ± standard error.

Plasma corticosterone concentrations were assessed in response to 20 minutes of restraint stress. Both groups of animals exhibited a robust increase in CORT due to restraint (Figure 2E, GLMM, Z = 6.9, p < 0.0001) while there was also a main effect of condition (GLMM, Z = 2.175, p = 0.03) with MS180 animals having overall higher plasma CORT concentrations across both stress and baseline periods. Additionally, female animals exhibited higher overall levels of CORT (GLMM, Z = −3.89, p < 0.0001). However, there was no interaction between maternal separation and restraint stress. As another measure of stress responsiveness, activation of the PVN following restraint stress was assessed through cFos immunohistochemistry (Figure 2F). When the number of cFos^+^ cells was assessed, a trend towards an interaction between condition and sex was observed (Figure 2G, GLMM, Z = 1.74, p = 0.083). When this was investigated it became apparent that male MS180 animals showed increased cFos expression compared to controls (GLMM, Z = 2.26, p = 0.024) while there was no difference in the female cohort. A main effect of sex was also observed overall with females showing higher cFos expression in response to restraint stress than males (GLMM, Z = −3.39, p = 0.0007).

Finally, neurogenesis in the dentate gyrus was examined as a biomarker of the ELS phenotype using BrdU immunohistochemistry (Figure 2H). There was no difference between control and MS180 groups in the number of BrdU+ cells in the SGZ of the DG (Figure 2I). There was additionally no effect of sex, although the total number of BrdU+ cells was higher in the VH as opposed to the DH (GLMM, Z = −6.92, p < 0.0001).

### 3.2 MS180 animals show increased NMDAR function compared to controls

To assess the relative function of AMPAR and NMDAR mediated CA1 pyramidal cell transmission AMPA/NMDA ratios were assessed in the SC and TA pathways of CA1 in control and MS180 animals (see Figure 3A for recording setup). MS180 animals overall had a lower AMPA/NMDA ratio compared to controls (Figure 3B, H and I, GLMM, Z = −3.16, p = 0.0016) with an additional interaction between sex and condition (GLMM, Z = −2.81, p = 0.005). When this was further investigated a clear effect of condition in females but not males was apparent (females: GLMM, Z = −2.31, p = 0.021). An overall trend for the TA pathway to have a lower AMPA/NMDA ratio was also observed (Figure 3J, GLMM, Z = −1.802, p = 0.072) with this manifesting as an interaction between region and pathway (GLMM, Z = 2.13, p = 0.03) whereby the TA pathway had a lower ratio in the DH (GLMM, Z = −2.04, p = 0.041) but not VH. This additional pathway analysis also elucidated an interaction between sex and condition in the DH with male MS180 animals only having a reduced AMPA/NMDA ratio in the DH (GLMM, Z = −2.50, p = 0.013).

**Figure 3.**
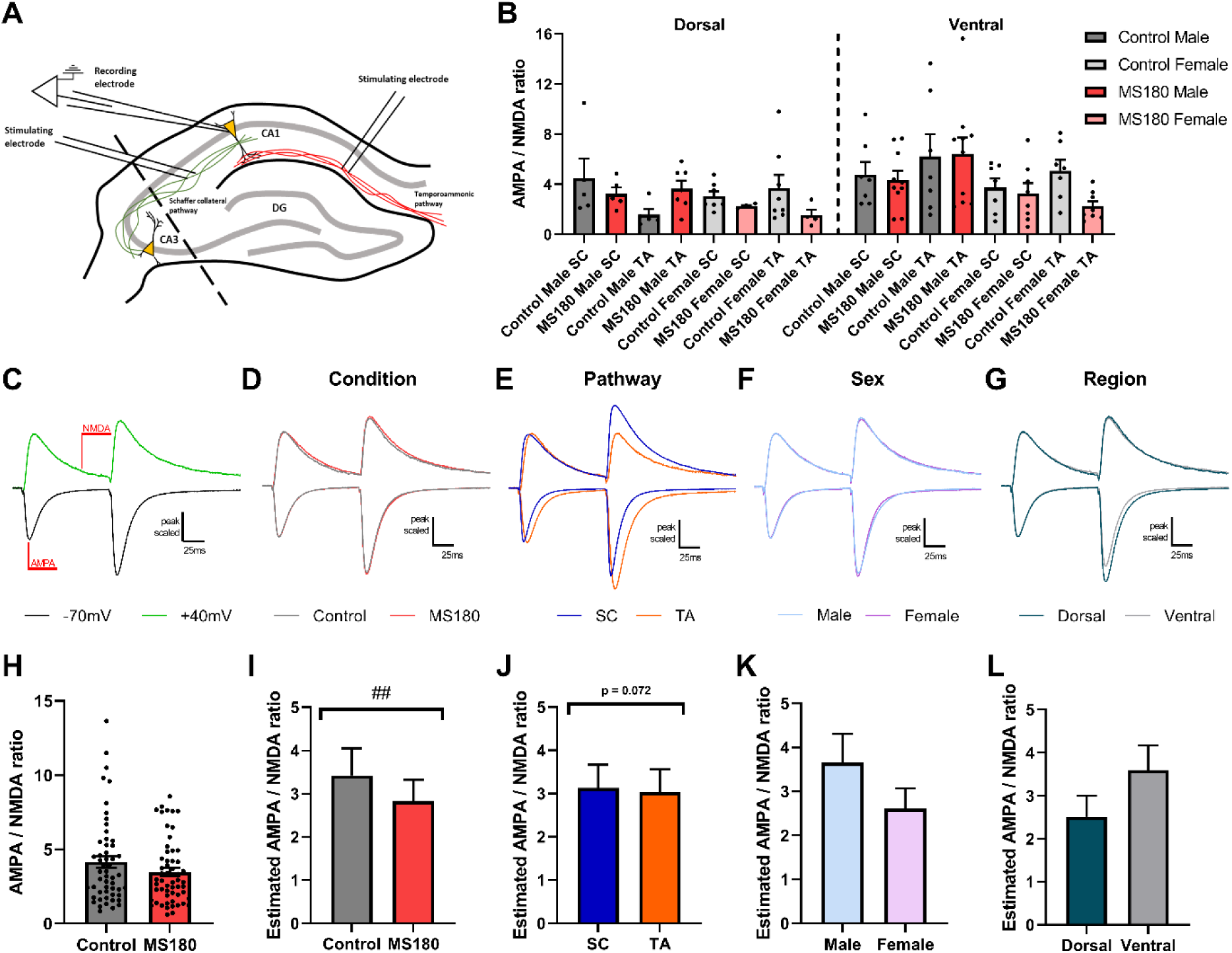
MS180 animals have a lower AMPA/NMDA ratio than controls. **(A)** Diagram of experimental setup with cells being recorded in CA1 and stimulated in SC and TA pathways. **(B)** AMPAR/NMDAR ratio. **(C)** Example trace showing where AMPAR and NMDAR mediated EPSC components were measured from. **(D-G)** Peak scaled average traces for the factors: condition, pathway, sex, and region. Downward deflecting traces indicate recordings at −70mV while upwardly deflecting traces indicate recordings from +40mV. **(H)** Example graph showing AMPA/NMDA ratio when all data is pooled by condition. **(I-L)** Estimated marginal means for condition, pathway, sex, and region whereby the effect of changing only the factor of interest is examined. N = 53 cells (26 control and 27 MS180) from 37 animals (18 control, 19 MS180) and marginal means are shown as mean ± 95% CI.

To isolate the effects of a changed AMPA/NMDA ratio to either AMPAR or NMDAR mediated transmission, miniEPSCs were recorded from CA1 pyramidal neurones (see Figure 4A for recording setup and Figure 4B/4C for example and average traces respectively). miniEPSCs were analysed as averaged events per cell to reduce any potential pseudoreplication. There was no difference in the amplitude (Figure 4D), area (Figure 4E) nor inter-event interval (Figure 4F) between control and MS180 animals. There was additionally no change in paired pulse ratio between control and MS180 animals (Figure 4G, 4H) as an indicator of changes in presynaptic function.

**Figure 4.**
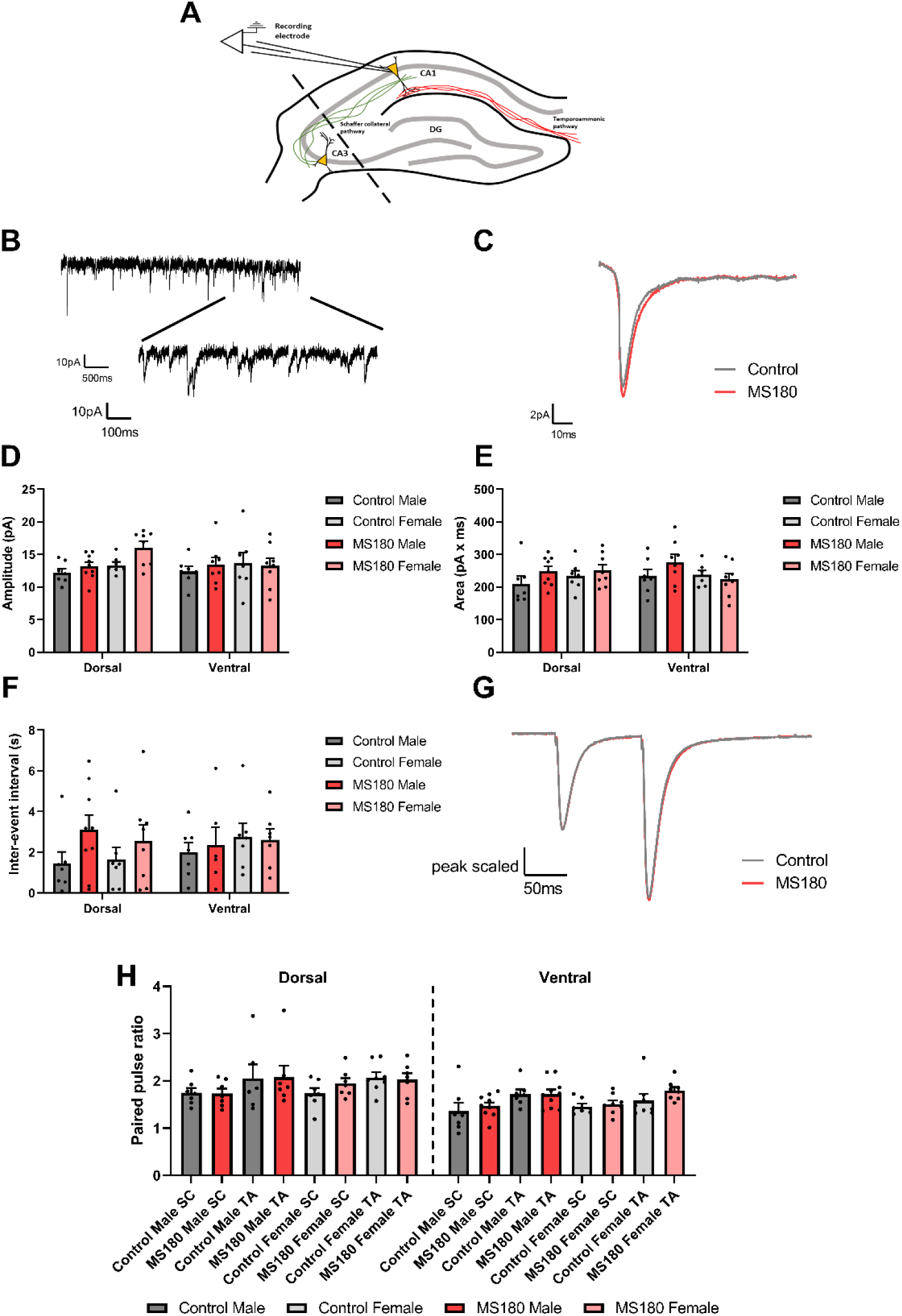
MS180 animals show no change in miniEPSC parameters. **(A)** Experimental setup diagram with cells recorded from in CA1 with no stimulation. **(B)** Example trace showing multiple miniEPSC events. **(C)** Average miniEPSC trace for control and MS180 animals. MiniEPSC amplitude **(D),** area **(E)** and inter-event interval **(F)** shown as per-cell averages. **(G, H)** Paired pulse stimulation average traces and per-cell average paired pulse ratio respectively. N = 61 cells (28 control and 33 MS180) from 19 animals (9 control, 10 MS180). Data shown as mean ± standard error.

### 3.3 Long term potentiation in MS180 and control animals

To ascertain if increased NMDAR function would translate into altered synaptic plasticity properties, theta burst stimulation induced LTP was studied in control and MS180 animals in both SC and TA pathways (see Figures 5A, 5B for overview). Due to a low n-number caused by Covid-19 related disruption all experiments whenceforth were analysed as control vs MS180 without interrogating the impact of animal sex or litter. All analysis was therefore conducted as either ANOVAs or T-tests as appropriate. Both groups exhibited robust short-term potentiation in Schaffer collateral but not Temporoammonic synapses (Figures 5C, 5D, 5E, 2-way ANOVA, main effect of pathway: F_1.76, 77.44_ = 14.88, p<0.0001). When long term potentiation was analysed, there was again a main effect of pathway (Figures 5C, 5D, 5F, 2-way ANOVA, main effect of pathway: F_1.29, 16.8_ = 7.24, p = 0.011) and no effect of condition (p=0.77). However, in post-hoc tests LTP was observed in control SC synapses (Dunnet’s multiple comparison, q_7_ = 3.3, p = 0.023) but not MS180 SC synapses (p=0.18).

**Figure 5.**
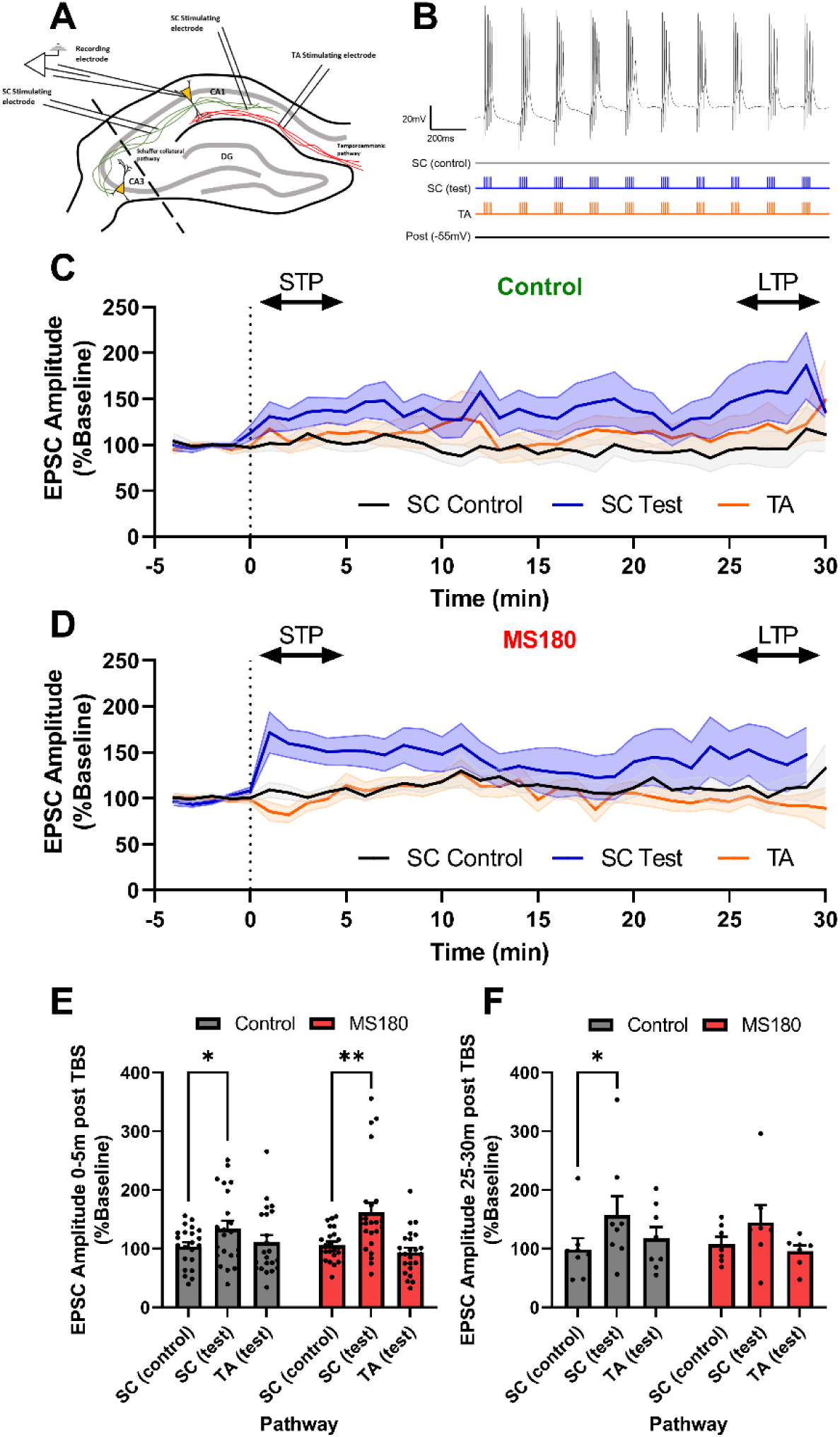
No change in short- or long-term potentiation between MS180 and control animals. **(A)** Diagram showing experimental setup with cells recorded in CA1 and two stimulation electrodes in stratum radiatum alongside a single electrode in stratum lacunosum-moleculare. **(B)** The theta burst stimulation LTP induction protocol showing an example trace resulting from it and a diagrammatic representation of the stimulation pattern. **(C, D)** Minute averages of normalised EPSC amplitude for control and MS180 animals respectively. LTP was induced at minute zero and STP/LTP markers represent where these measurements were taken from. **(E, F)** STP (0-5 mins post TBS) and LTP (25-30 mins post TBS) respectively split by pathway and condition. STP: n = 46 cells (22 control and 24 MS180); LTP: n=15 cells (8 control, 7 MS180) – both from 20 animals (10 control, 10 MS180). Data shown as mean ± standard error.

Data from LTP induction were analysed to understand if differential excitability during the TBS protocol could be present between control and MS180 animals that might affect their ability to express LTP (see Figure 6A for overview). There was no difference between control and MS180 animals observed when the total number of action potentials (Figure 6B), area under the theta burst (Figure 6C) nor the decay area following each theta burst (Figure 6D, a measure of NMDAR activation) were analysed. There were additionally no changes between conditions in input resistance (Figure 6E) that might explain any changes in NMDAR function.

**Figure 6.**
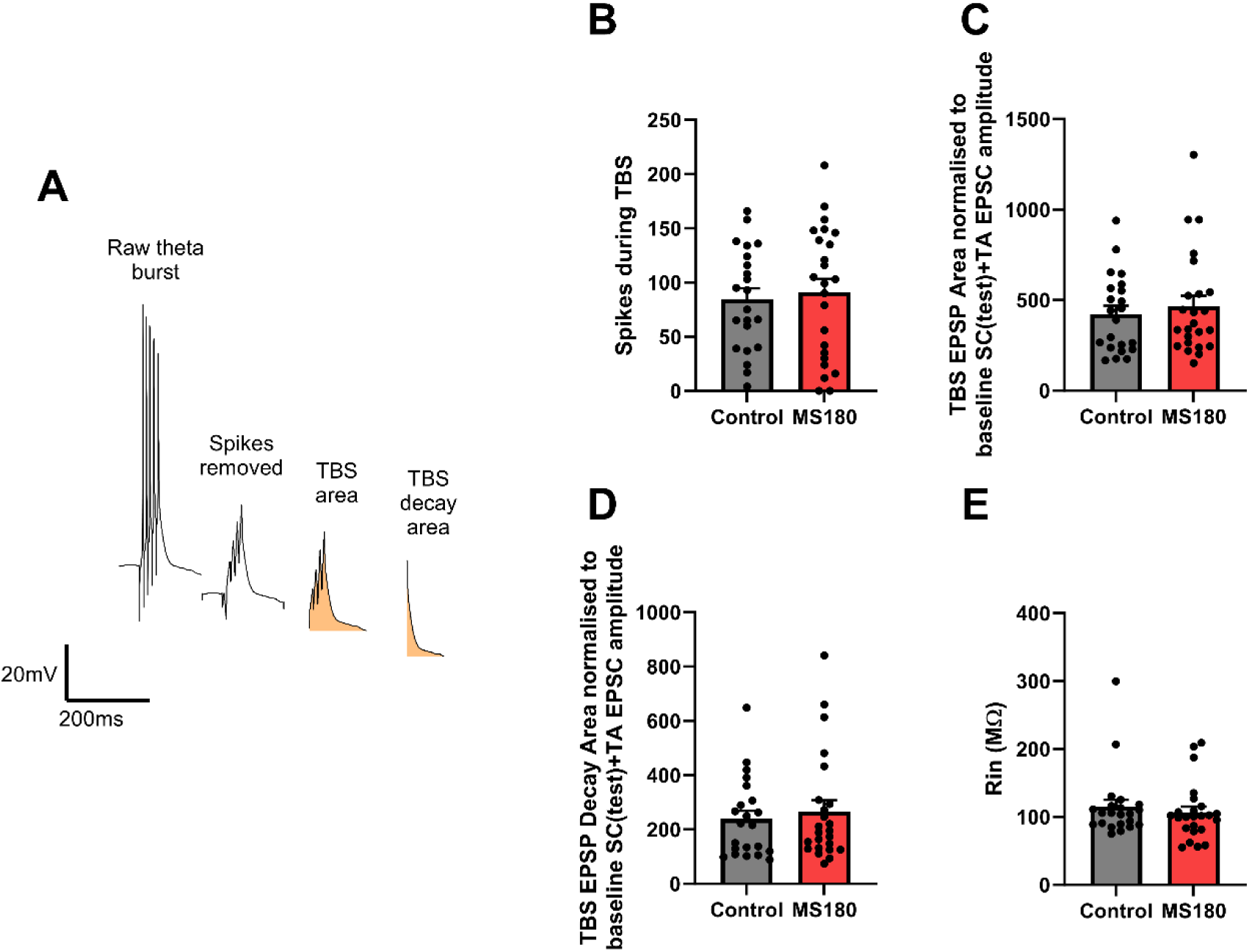
No effect of maternal separation upon theta burst parameters or input resistance. **(A)** Example trace showing how output measures were created. **(B)** Total number of spikes counted during the theta bursts. **(C, D)** Total EPSP area and area from the decay phase respectively from the LTP induction phase. **(E)** Input resistance (R_in_). N = 46 cells (22 control and 24 MS180) from 20 animals (10 control, 10 MS180). Data shown as mean ± standard error.

## 4 Discussion

### 4.1 Model Validation

Initially a battery of experiments were carried out to establish that animals displayed a phenotype consistent with that previously seen in ELS animals. MS180 animals displayed increased anxiety as evidenced by an increased latency to feed in addition to increased thigmotactic behaviour in the NSFT. This matches well both with previous data from the NSFT (Bonapersona et al., 2019; Stuart et al., 2019). The increase in proposed thigmotactic behaviour was interesting considering that a recent meta-analysis concluded that maternal separation did not lead to increased anxiety in the open field test (Wang et al., 2020). Interestingly the effects of maternal separation upon anxiety may be test specific with the same study reporting significant effects of MS upon behaviour in the elevated plus maze. In the present study there was also no difference between male and female rats in terms of anxiety which compares to reports that ELS leads to greater anxiety in male rather than female animals (Bonapersona et al., 2019). As reported previously (Shalev and Kafkafi, 2002; Stuart et al., 2019), MS180 rats in the present study did not show changes in sucrose preference indicating no changes in hedonic behaviour.

Although during the restraint stress CORT experiment a main effect of ELS was observed this was not consistent with previous studies suggesting that MS180 animals should have potentiated responses to stress but no changes at baseline (Aisa et al., 2007; Plotsky and Meaney, 1993; Stuart et al., 2019). However, as seen with humans following ELS (Bunea et al., 2017; Fogelman and Canli, 2018) there is considerable heterogeneity in the HPA axis consequences of maternal separation. Other studies have reported decreased ACTH secretion (Daniels et al., 2004; Marais et al., 2008) and decreased CORT release (Roman et al., 2006) following restraint stress in MS180 animals relative to controls. Interestingly it has been observed that maternal care and maternal separation act independently upon HPA axis outcomes with high levels of maternal care able to compensate for extended maternal separations (Macrì et al., 2008). Differences in maternal care being able to compensate for maternal separation may be the cause of much heterogeneity between studies with Long-Evans mothers reported to show an increased quality of maternal care relative to Wistar and Sprague-Dawley mothers (McIver and Jeffrey, 1967).

Male MS180 rats appeared to exhibit increased cFos activation of neurones in the PVN following restraint stress in a result matching previous observations (Sanders and Anticevic, 2007). However for males to have higher restraint stress PVN cFos but not increased CORT suggests downstream changes in the HPA axis such as decreased ACTH release (Daniels et al., 2004; Marais et al., 2008) or altered CORT feedback regulation (van Oers et al., 1998). However, many of the PVN neurones release oxytocin as opposed to CRH (Nishioka et al., 1998) with it not being possible to know which population the increase in activation resulted from.

There was no difference between control and MS180 animals with respect to dentate gyrus neurogenesis utilising BrdU labelling. This contrasts with previous studies that reported robust decreases in proliferation (Mirescu et al., 2004; Stuart et al., 2019). One potential reason for differences between studies is animal age with animals in the discussed studies being between PND60 and PND70 while animals in the present study were aged PND112-123. A previous study investigating neurogenesis across the life course reported that proliferation as measured by Ki67 staining at PND120 is between 50 – 100% lower than at PND70 (Epp et al., 2009). This could mean that by PND123 neurogenesis had decreased to a degree where detecting further decreases would be challenging.

Overall, while not completely matching previous data, these validation experiments give enough confidence that a phenotype matching that previously seen in other ELS models was created to allow interpretation of electrophysiological data.

### 4.2 NMDAR Function and LTP

Female MS180 animals showed a reduced AMPAR/NMDAR ratio while showing no changes in miniEPSC amplitude. Due to the fact that miniEPSCs recorded at −70mV are AMPA dependent (Kato et al., 2007) this indicates the MS180 females have increased NMDAR function. This is in contrast to Pillai et al., 2018 who reported decreased NMDA receptor function relative to AMPA. However this was carried out in male mice using a limited nesting and bedding material model (LNBM) of ELS which appears to generate a different phenotype to the rat MS180 model focussed more upon spatial learning and memory deficits with little evidence for an emotional deficit (Bath et al., 2017; Kanatsou et al., 2017; Rice et al., 2008; Wilkinson, 2021).

Brunson et al., 2005 also tried to measure NMDA function by assessing EPSC decay at membrane voltages between −80mV and 0mV using the LNBM model, albeit in rats. They did not observe any changes in NMDA function between ELS and control animals, however again only males were assessed. It should be noted that an alternative explanation for these results may be increased numbers of silent synapses in female MS180 animals (Kullmann, 1994). The lack of changes observed in the present study in AMPA receptor function are also in contrast to previous work which observed decreased GluR1 and GluR2 mRNA expression in rats that were maternally separated for 360 minutes per day (Pickering et al., 2006).

Before discussing further, it is worth reiterating that data regarding LTP in MS180 animals is underpowered due to limitations imposed by the effects of Covid-19 and therefore needs interpreting cautiously. There was no difference in LTP nor STP between MS180 and control animals with the lack of SC LTP in ELS animals likely due to the low statistical power in the experiment. Interestingly it should be noted that in the present study LTP was assessed in naïve animals that had no other manipulations. Previous data from maternally deprived animals in the DG reported no differences between controls and ELS animals at baseline but greater LTP following stress exposure via CORT injection (Oomen et al., 2010). There was a general failure to induce LTP in the TA pathway in the current study which may be due to a lower baseline EPSC amplitude meaning that not enough depolarisation was generated during LTP induction to generate LTP. As previously discussed other studies have observed increased LTP following ELS (Derks et al., 2017) which matches that which would be predicted by the increased NMDA function seen in ELS females. The fact that there was no effect of hippocampal region upon LTP is interesting in itself and matches with Kouvaros and Papatheodoropoulos, 2016 who reported no difference in LTP magnitude between regions but an increased ability for induction and stability of LTP in the DH as opposed to VH. There was also no effect of early life stress upon spiking or depolarisation during the theta burst LTP induction protocol suggesting both groups can generate robust NMDAR mediated responses to TBS stimulation.

There are interesting parallels to the results presented by Griesius et al., 2022 who also found an increase in NMDAR function in a psychiatric risk model (DLG2 heterozygous knockout) with no change in AMPAR function. They reported that the effects on LTP are dependent upon the stimulation protocol with associative LTP being impaired but not with LTP induced by pairing between the pathway inputs. Additionally they deduced that increased NMDAR function did not lead concurrent increase in LTP due to a concomitant decrease in input resistance. This implies that the effects of MS180 upon LTP may be stimulation protocol dependent and that the increased NMDAR observed may be prevented from increasing LTP through other mechanisms. Interestingly input resistance was unchanged between MS180 and control animals suggesting that another mechanism other than the one identified in Griesius et al., 2022 may be in play.

### 4.3 Conclusions

These data suggest that although the MS180 model did not produce fully congruent results with previous literature it was successful in generating an anxiety phenotype suggesting a believable foundation for investigating neural circuit changes. MS180 females showed a decreased AMPAR/NMDAR ratio but no change in miniEPSC amplitude implying an increase in NMDAR function. This did not translate into changes in LTP or STP nor dynamics following TBS. These findings warrant further investigation to complete this detailed investigation into the hippocampal consequences of ELS taking into respect hippocampal aspect, sex and pathway in a way not previously done before.

## 5 Data Availability

All data underlying this manuscript is available at: www.osf.io/mbhzk. The project contains the following underlying data:

- Raw data (raw experimental files and analysis scripts)
- AMPAR_NMDAR data (summary data of AMPAR/NMDAR experiment)
- miniEPSCs data (summary data of miniEPSC experiment)
- LTP data (summary data of LTP experiment)
- AMPAR_NMDAR average traces (average traces from AMPAR/NMDAR experiment)
- Validation and breeding data (combined data from validation experiments and information on animal breeding)

Data are available under the terms of the Creative Commons Attribution 4.0 International (CC BY 4.0)

## Acknowledgements

The authors would like to acknowledge Simonas Griesius for assistance in breeding of animals, Daryl Purawijaya for aid in analysis of NSFT data followed by Julia Bartlett and Sarah Stuart for helping conduct the restraint stress corticosterone and BrdU neurogenesis experiments. The authors would also like to thank Michelle Taylor for advice regarding statistical analysis and the Wolfson Bioimaging Facility for support and assistance with imaging. Finally the authors would like to extend thanks to all members of the Robinson and Mellor laboratories alongside members of the animal services unit at the University of Bristol.

